# Inhibition of a critical malaria host-pathogen interaction by a computationally designed inhibitor targeting *Plasmodium vivax* DBP

**DOI:** 10.1101/2022.02.05.479252

**Authors:** Autumn R Tobin, Rachel Crow, Darya V. Urusova, Jason C. Klima, Niraj H. Tolia, Eva-Maria Strauch

## Abstract

Malaria is a substantial global health burden with 229 million cases in 2019 and 450,000 deaths annually. *Plasmodium vivax* is the most widespread malaria-causing parasite putting 2.5 billion people at risk of infection. *P. vivax* has a dormant liver stage and therefore can exist for long periods undetected. Its blood-stage can cause severe reactions and hospitalization. Few treatment and detection options are available for this pathogen. To address this need, we developed nanomolar inhibitor that could serve as a therapeutic and a diagnostic. A unique characteristic of *P. vivax* is that it depends on the Duffy antigen/Receptor for chemokines (DARC) on the surface of host red blood cells for invasion. *P. vivax* employs the Duffy binding protein (DBP) to bind to DARC. We first *de novo* designed a three helical bundle scaffolding database which was screened via protease digestions for stability. Protease-resistant scaffolds highlighted thresholds for stability, which we utilized for selecting DARC mimetics that we subsequentially designed through grafting and redesign of these scaffolds. The optimized design small helical protein disrupts the DBP:DARC interaction. The inhibitor blocks the receptor binding site on DBP and thus forms a strong foundation for a therapeutic that will inhibit reticulocyte infection and prevent the pathogenesis of *P. vivax* malaria.

**Teaser:** *De novo* designed proteins present a new alternative for the development of therapeutics. They can be small, highly stable and easily manufactured. Here we designed a potential new therapeutic to inhibit entry of *Plasmodium vivax* into red blood cells by interfering its interactions of surface displayed DBP molecules with the host receptor DARC.

## Introduction

Malaria is predicted to infect up to 2.5 billion by the year 2050. *Plasmodium vivax* has been the second largest cause of human malaria around the world after *Plasmodium falciparum*. However, substantial increases in malaria incidence occurred between 2014 and 2016, with *Plasmodium vivax* representing 64% of all malaria cases(*1*). Most strategies deployed to eliminate malaria primarily target falciparum malaria and are less effective in controlling vivax malaria(*2*). Basic research on *P. vivax* has been hampered by lack of continuous *in vitro* culture system. Furthermore, *Plasmodium vivax* poses unique challenges to control strategies because of its ability to cause relapses(*3*); it can remain dormant in the liver for years before entering its blood stage form in which several clinical symptoms can occur. Primaquine is the drug currently used to eliminate hypnozoites from the liver. However, adverse effects associated with primaquine have been reported, particularly in patients who have a severe deficiency of the enzyme glucose-6-phosphate dehydrogenase (G6PD), in whom the drug triggers hemolysis. Therefore, primaquine cannot be administered to pregnant women or children because of the risk that the patient might have this enzyme deficiency and new treatment options are needed.

During the life cycles of *P. vivax*, different surface proteins are displayed at various stages of the parasite. During its blood stage cycle, the entry of host cells by the parasite is mediated by the Duffy Binding Protein (DBP) (*4–12*), a member of the Erythrocyte Binding-Like (EBL) invasion protein family (*13–15*). DBP binds to the Duffy Antigen Receptor for Chemokines (DARC) on host reticulocytes using a conserved cysteine-rich Duffy Binding-Like (DBL) domain known as region II (DBP-II) (*4–11*). DBP-II binds DARC via receptor-induced ligand dimerization, sandwiching DARC residues 19–30 between two DBP-II molecules (*9, 10*). Receptor binding and dimerization are required to engage DARC (Fig.1) (*9, 10*). The stepwise binding mechanism improves the affinity and avidity of the DBP-II:DARC interaction leading to nanomolar binding and stable complex formation (*10*).

**Figure 1.**
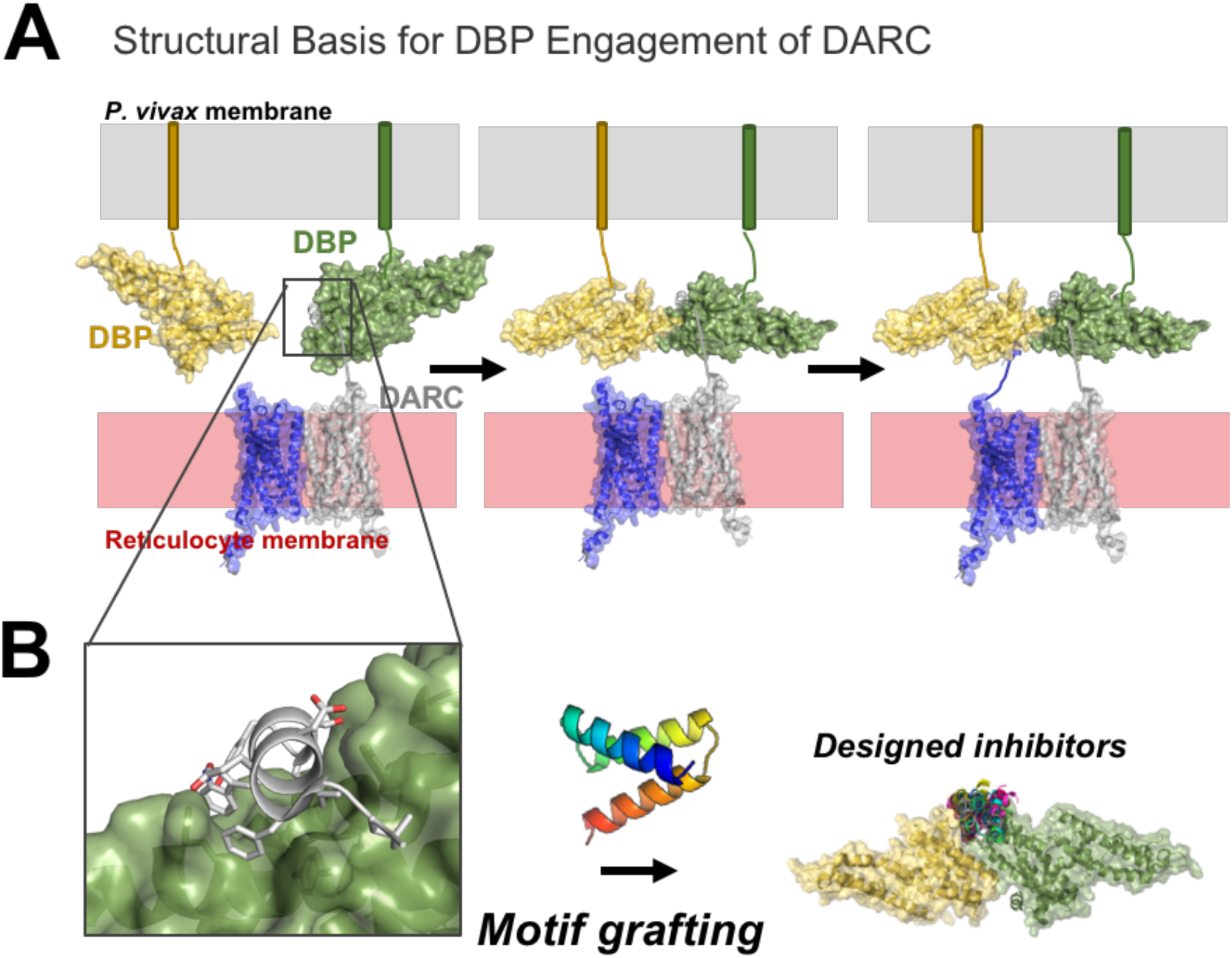
(A) Binding mode of DARC with DBP-II when membrane embedded. DARC is based on the Alphafold homology model AF-Q16570-F1 (Uniprot). (B) Design of receptor mimicries and their interference of DBP binding and dimerization.

Antibodies from immune individuals with naturally acquired immunity that block the DARC receptor binding and/or dimerization neutralize *P. vivax* (*9, 10, 16–18*). A single human monoclonal antibody that inhibits the DBP:DARC interaction is a potent neutralizer of *P. vivax* invasion of human red cells demonstrating that disruption of the DBP:DARC interaction is sufficient to prevent *P. vivax* invasion of host cells (*18*). DBP-II is a promising vaccine candidate for *P. vivax* malaria given the critical role of the DBP:DARC interation in *P. vivax* host cell invasion (*9–11, 19–25*) In addition to its vaccine potential, DBP-II presents an ideal target molecule for a protein-based inhibition, and a therapeutic that can block both receptor binding as well as interfere with the dimerization is highly desired.

To develop a potent inhibitor, we started by designing a highly stable, helical scaffold library, and used this library to develop a series of receptor mimicries that inhibit the interaction of DBP-II with its host cell receptor. We designed and experimentally evaluated 4,000 helical bundles of the size of 48 – 68 residues long (Fig. S1) and then monitored stability by a direct protease-digestion assay using yeast surface display. Proteins that survived protease-digestion as identified by next-generation sequencing were used as starting material for the design of the receptor mimicries. Surprisingly, this stringent selection showed clear cut-offs for scoring terms of these helices (Fig. S2) and resulted in a set of highly stable scaffold proteins. The results highlight stability thresholds for these *de novo* designed helical bundles, identifying selection criteria for the future design of different topologies as well as the re-design of the original scaffolds produced here. Proteins were redesigned by grafting the core interactions of DARC with DBP-II on the initial designs. Our lead candidate binds with single digit nanomolar affinity to the receptor binding site of DBP. The angle of binding of the lead candidate to DBP is such that it would also prevent dimerization of DBP. Cell-based assays demonstrate the lead candidate potently inhibits the binding of DBP to DARC on red blood cells with an IC50 of 72.5 nM, and these results form a strong foundation for the development of a potent neutralizing therapeutic reagent.

## Results

A protein-based inhibitor that has to serve both as a potential therapeutic as well as diagnostic for treating and diagnosing a *P. vivax* malaria infection would have to be suitably stable to avoid degradation within the body, and posses characteristics that would enable long-term storage at temperatures suitable for deployment in the field. Furthermore, high expression of protein scaffolds are also desired to ensure affordable production costs. Lastly, a designed protein inhibitor must bind to DBP and bury a large enough surface area to mimic the receptor interaction while also sterically blocking the dimer interface of DBP-II.

We therefore first designed a “scaffold” library of all helical three helical bundles with varying sizes and defined loop angles. This library is comprised of larger proteins than the previously reported mini-proteins that were 42 residues long and less(*26*) as the binding interface in DBP-II we wanted to graft was large and contained several hydrophobic residues(*10*). We reasoned that a scaffold presenting this binding interface would need to be larger than previously reported mini-proteins(*26*) and the hydrophobic residues on the surface should be fewer in number than the number in the core of the protein to avoid misfolding or instabilities. Our newly generated helical scaffold proteins range from 48 to 68 amino acid residues. They provide a starting point for either computational redesign or as a starting scaffold for more traditional approaches for protein engineering using library generation and selections.

### De novo design of scaffold protein library

For the *de novo* design of proteins using the macromolecular software suite Rosetta(*27*), a blueprint file is needed to describe a single topology with fixed lengths of all helical residues and specific *phi* and *psi* angles for the connecting loops. The backbone angles for the loop residues are defined using the coarse-grained ABEGO terminology (*28*) which describes the *phi/psi* angle combinations of the areas populated in natural proteins and summarized by the Ramachandran plot. To exhaust the conformational space possible, we allowed all permutations that had either 14, 15 or 16 residue long helices or in a second set 17, 18 and 19 residues together with one of the 4 possible loops descriptions (BB, GBB, GB, GABB) for the two loops. We kept the lengths of the helical elements within each set at described size ranges to avoid producing overly uneven sized helices which could potentially result in unstable overhangs. This resulted in 432 possible blueprints to fold a three helical bundle topology. Since not all of these permutations will result in reasonable topologies, we allowed *in silico* folding through fragment insertion(*29*) following sequence design for a short defined time period to identify reasonable topology descriptions; resulting decoys were filtered based on backbone geometry, similarity to existing four-mers in the PDB, helix bending, and packing (see Methods and SI). Blueprint descriptions that produced large numbers of models were labeled as promising and scaled up in their production time; as expected not all of them produced an output (Fig. 2A).

**Figure 2.**
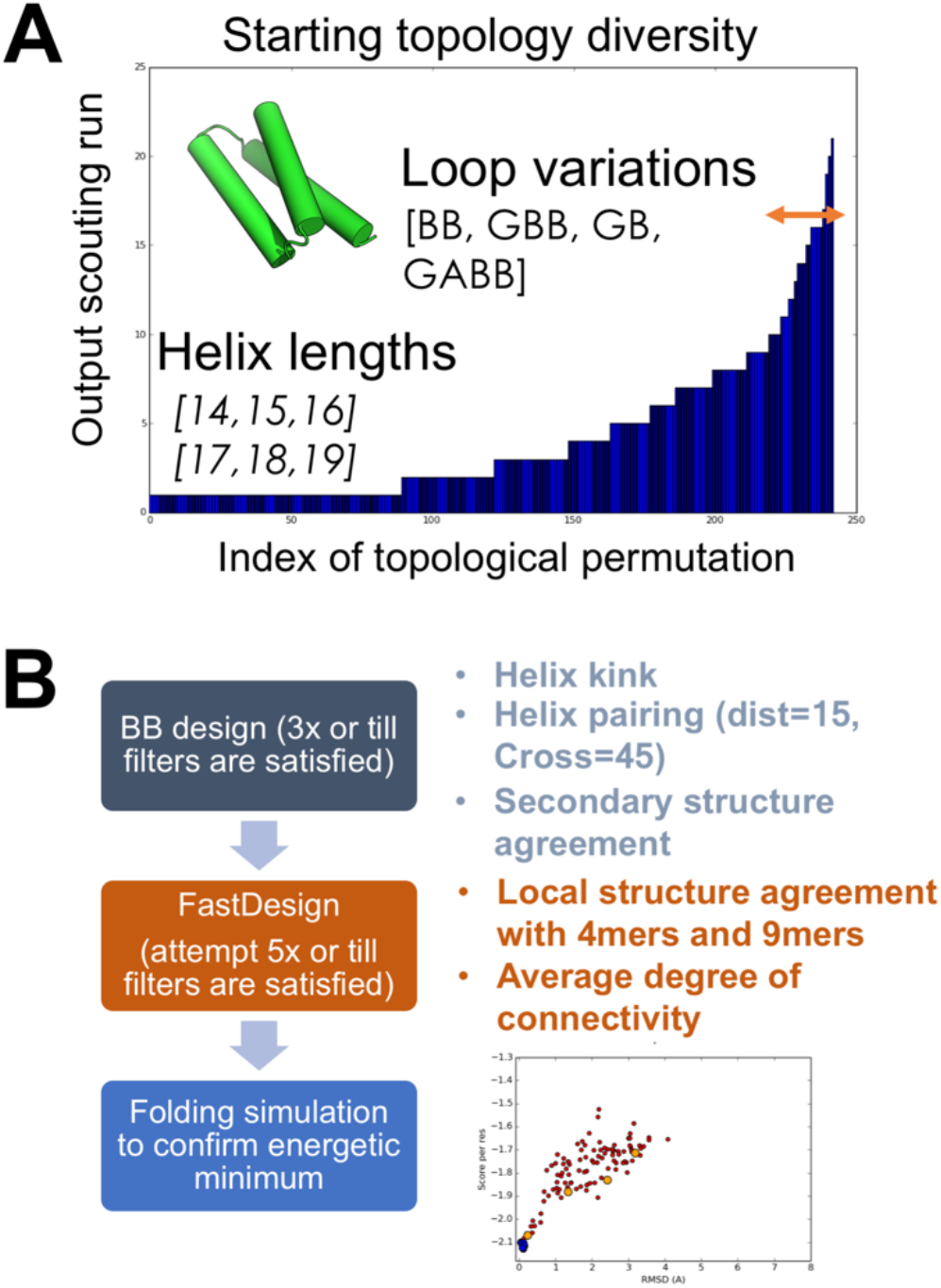
(A) Histogram illustrating which topology permutation can be successfully made after an initial time limited design trajectory. (B) Overview of the design workflow for the new helical scaffold proteins.

### Stability selection using yeast surface display

A DNA sequence of the amino acid sequence of the newly designed proteins was then generated using DNAworks(*30, 31*), and sequences were ordered in form of a 230 bp long oligonucleotide pool together which also contained adapters for cloning into a yeast surface expression plasmid as reported before(*32*). Designs were expressed on the surface of yeast and subjected to three different concentrations of proteases (a mix of both chymotrypsin and trypsin, Fig. 3A, B and Method section). After two rounds of sorting, plasmids were extracted from the sorted pools and starting pool, then subjected to next-generation sequencing to obtain enrichment values. Designs with increasing enrichment comparing across the three different concentrations of proteases were considered as stable. While this variation of a screen for folded proteins is less quantitative as a previously reported method in which stability scores can be derived based on computed EC50 values of independent protease titrations (keeping trypsin and chymotrypsin separate); this variation allows for a more efficient and direct way to identify folded and stable proteins. We compared basic Rosetta score terms of successful designs and noticed a minimum threshold for several of the parameters, particularly relevant to the smaller helical bundle set (Fig. 3C, S2). These cutoff values can be used for either redesigning these scaffold proteins or for developing scaffold sets. Out of a total of 4,000 screened sequences, 800 design models were identified to be highly protease resistant. These 800 were then used to graft the DARC binding site onto using Rosetta’s motif grafting protocol(*33*).

**Figure 3.**
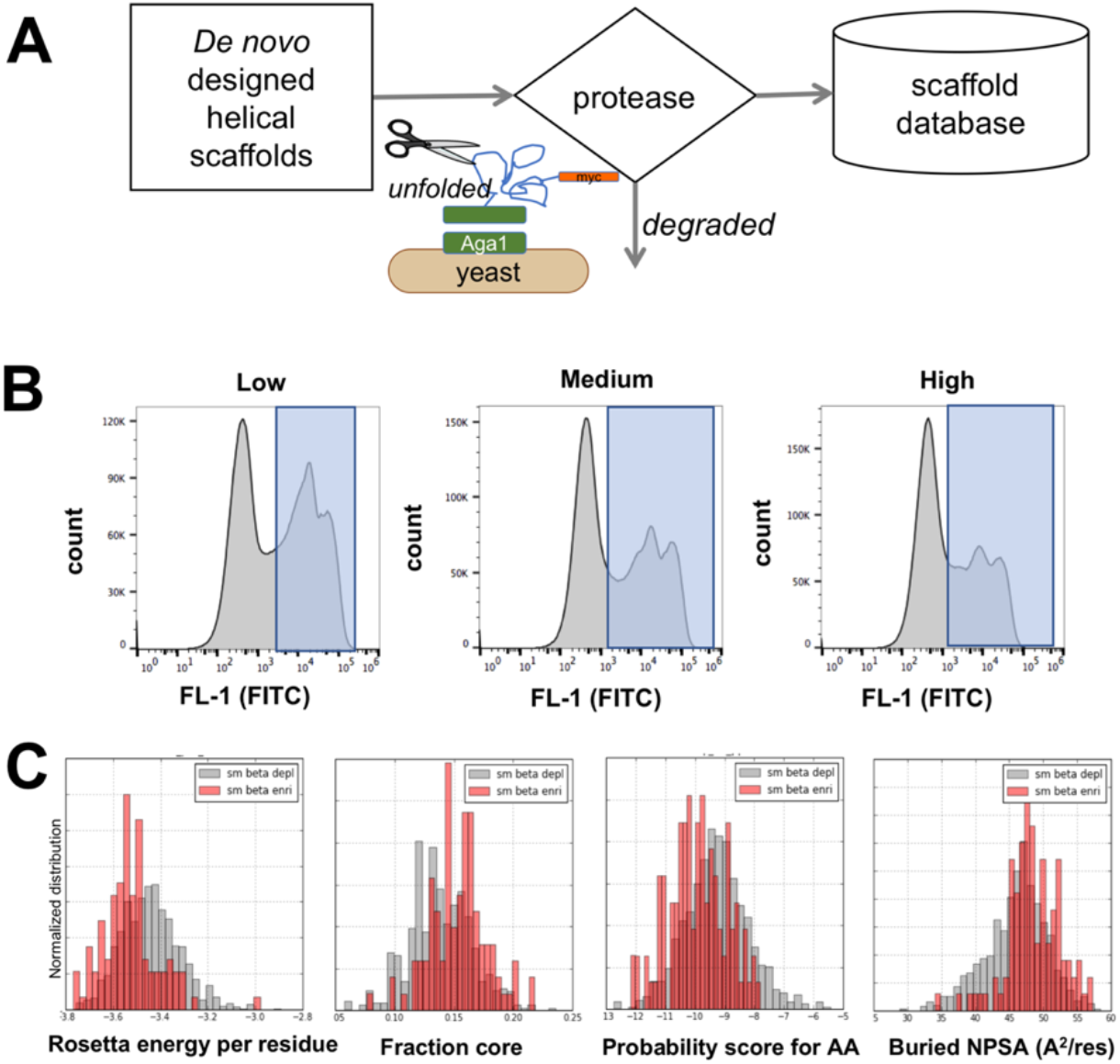
Summary of initial yeast surface display screen. (A) Scheme selecting of folded designed proteins using yeast surface display combined with protease digestions. (B) Data of selection of the second round of selection at the 3 different protease conditions. (C) Identified thresholds for our designed helical bundles.

### Grafting of the DARC binding interface and characterization

We kept hotspot residues as well as additional interface residues (22,23,25,26) of the core interacting helix unchanged but allowed all other residues to vary. Furthermore, different lengths of the interface helix were sampled (Methods). Designs with low-binding energy were then evaluated for their folding capability *in silico* using *ab initio* structure prediction (*34*). Only designs that had clear energy funnels with a minimum close to the designed conformation were ordered (Fig. S3).

Sixteen designs were synthesized as gBlock (IDT DNA) and tested *via* yeast surface display for binding to 10 μM purified, biotinylated DBP-II (Fig. S4). All designs showed a distinct binding signal at these high concentrations. To ensure that the designs were folded and highly stable, we evaluated their expression in *E. coli*. Twelve designs were successfully cloned in an expression vector and 8 designs expressed soluble protein. All soluble designs showed a dominant, monodisperse peak at the expected elution volume by size-exclusion chromatography. Circular dichroism spectrometry established that the designs were indeed folded and helical, and recorded distinct alpha-helical pattern for 6 of the 8 purified *de novo* designed proteins (Fig. 4). All folded designs had a melting point above 95°C and even at that high temperature, a clear helical profile was recorded except for Dbb10 (Fig. 4C).

**Figure 4.**
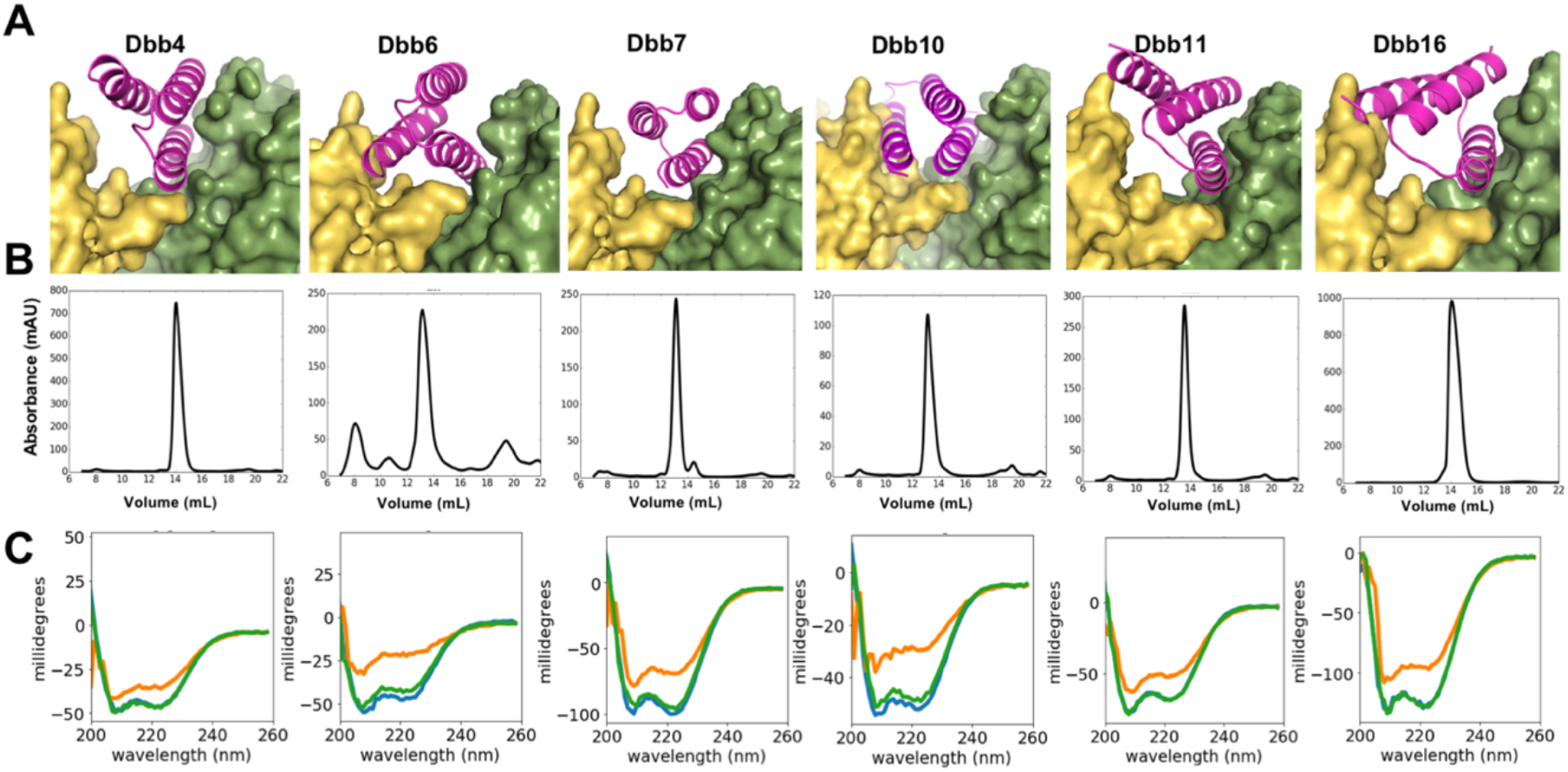
DARC mimetics. (A) Model of the designed protein occupying the DBP-DARC interaction site. DBP chains are colored in yellow and green, whereas the design is colored in pink. (B) Size-exclusion chromatography monitored at 280 nm absorption wavelength of soluble DBP inhibitors after Ni-NTA purification from E. coli expression. (C) Far-ultraviolet circular dichroism at 25° C (blue), 95° C (orange) and at 25° C after heating it up to 95° C (green).

### Library screening

To optimize binding while maintaining high stability, we generated a library of all possible point mutations of Dbb4, Dbb7, Dbb10 and Dbb16 and selected for binding using fluorescent activated cell sorting (FACS). However, Dbb16 appeared to be the tightest binder, so we focused on this design. The site-saturation mutagenesis (SSM) libraries was screened for binding to 250 nM and 1 μM of biotinylated DBP-II. To determine enrichment values, plasmids from starting pool and selected populations were extracted and frequencies determined through next-generation sequencing using a NextSeq platform (Fig. 5).

**Figure 5.**
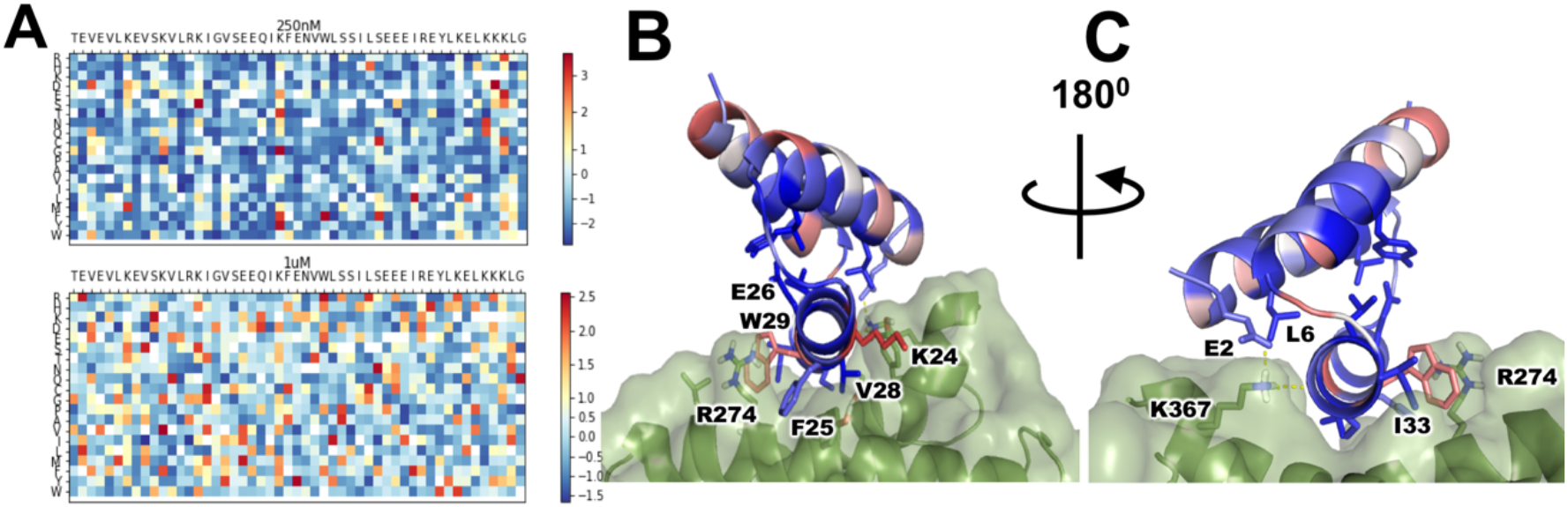
SSM library results of the lead candidate Dbb16. (A) Heat maps of enrichment values at 250 nM DBP-II and 1 μM DBP-II. Red indicates improvement of binding. (B) Model of Dbb16 colored by Shannon entropy computed from the enrichments; blue colors represent more conserved residues. (C) Model of bound Dbb16 turned by 180 degrees.

We computed the Shannon Entropy of its enrichment values to highlight conserved residues as well as positions that could benefit from optimization (Fig. 5). As expected, the core contacts F25 and V28 were highly conserved, as was the salt bridge between E26 and R274 of DBP-II. Interestingly, a phenylalanine was preferred instead of tryptophan at position 29. Also, K24 appeared to be suboptimal and more hydrophobic contact or a tyrosine was preferred, contradicting the Rosetta based designed residue at this position. Core residues found in Dbb16 were conserved indicating optimal packing of the small protein. Additional contacts to the targeted DBP-II are established by E2 and L6 which were also conserved. To optimize binding, we generated a combinatorial library for Dbb16 based on the SSM enrichment data (Table S1); amino acid variations were incorporated through degenerative primers with optimized codons using SwiftLib (*35*) thereby minimizing additional amino acid identities to be encoded.

To ensure high stability, we performed the first two rounds of selection by first incubating displaying yeast cells with the highest concentrations of trypsin and chymotrypsin (as used previously for the screening of the initial *de novo* designed scaffold library). After the incubation with the protease cocktail, cells were incubated with 200 nM of biotinylated DBP-II (bDBP-II) for round 1 and 50 nM bDBP-II for round 2 before sorting. The third and fourth rounds were performed at 5 nM and 1 nM (Fig. 6A). Clones after 3 and 4 rounds of sorting were sequenced (Fig. 6B). While round 4 converged onto a few sequences, more diversity was seen for round 3.

**Figure 6.**
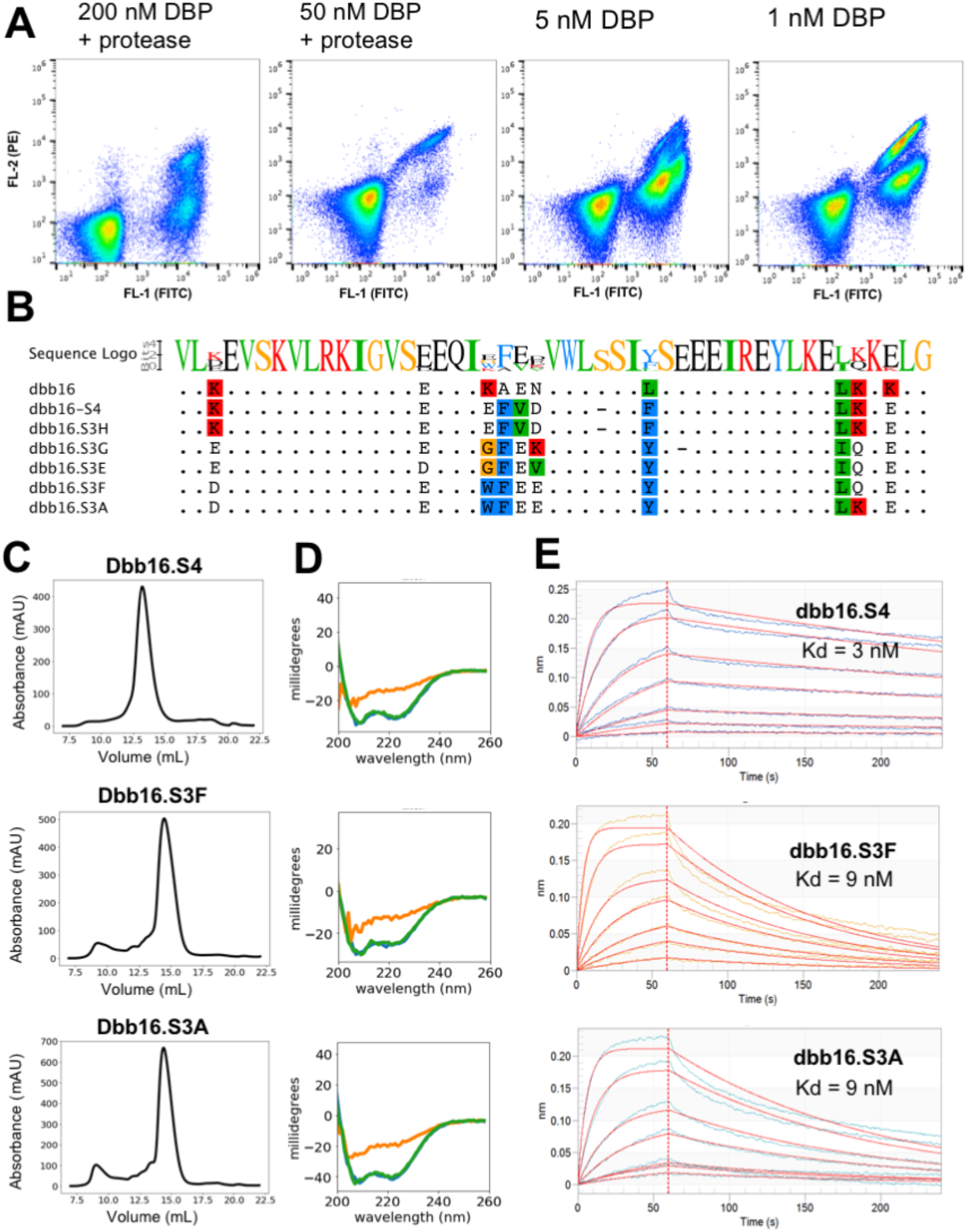
Library selections and biochemical characterization of Dbb16. (A) FACS rounds for affinity maturation of Dbb16. The concentration of bDBP-II was steadily descreased. Round 2 started by first incubating the displaying yeast cells with 0.25 μM trypsin and 0.05 μM chymotrypsin before washing and incubating with 50 nM bDBP-II. (B) Sequences of identified clones after 3 and 4 rounds of sorting. (C) SEC profiles monitored at 280 nm absorbtion wavelength of soluble identified variants after expression in E. coli. (D) Far-ultraviolet CD spectra of variants; 25° C (blue), 95° C (orange) and at 25° C after heating it up to 95° C (green). (E) Biolayer interferometry of variants binding to immobilized bDBP-II.

### Characterization of affinity maturated Dbb16 variants

Sequenced clones from rounds 3 and 4 were cloned and expressed solubly for further characterization. Proteins were purified as described for earlier variants and SEC showed monodispersed peaks at the monomeric fraction, except for Dbb16.S4. Interestingly, Dbb16.S4 had a deletion in one of its loops’ residues. *Ab initio* structure predictions using both Rosetta (*34*) and Alphafold2 (*36*) suggests that the protein variant has the same fold with a shorter loop and a small change at the end of the second helix leaving the interface and core intact (Fig. S5). As our affinity maturation process increased the hydrophobic content of the designed interface found in Dbb16.S4, it is likely to be the site for dimerization. We have noted before that highly hydrophobic interfaces of designed protein binders tend to dimerize at their designed interface. Structure predictions using Alhpafold2 provided a possible homodimeric model for Dbb16.S4 (Fig. S5). We then measured the secondary structures of the top 3 variants *via* CD and found all variants to be helical as before (Fig. 6D). Temperature melts showed that the affinity maturated variants do unfold at 95°C, but refold into their original conformation upon cooling down to room temperature. To confirm binding as soluble constructs, we measured binding using biolayer interferometry (BLI) and as expected, the converging sequence after 4 rounds of sorting bound the tightest (Fig. 6E) which bound three times tighter than the round 3 variants. Its improvements are derived from a slower off-rate.

### Inhibition of DBP binding to RBCs

A tremendous challenge to develop *P. vivax* treatment option is the fact that the parasite cannot be cultivated under laboratory conditions as well as to be grown in the actual Anopheles mosquito. As an approximation to see whether we could inhibit reticulocyte entry mediated through bDBP-II we utilized previously reported Fluorescence Activated Cell Sorting Assay that utilizes recombinant protein (*18*). Titrating our inhibitor Dbb16.S4 with the protein (0.01 nM - 3 μM interval of inhibitor’s concentration) we were able to determine an IC50 of 72.5 nM (Fig. 7A). The inhibitor fully occupies the DARC binding site and extends beyond the original contacts. Further, even though it is a small protein of 5.5 kDa, it should sterically block the dimerization of DBP as our model indicates (Fig. 7B).

**Figure 7.**
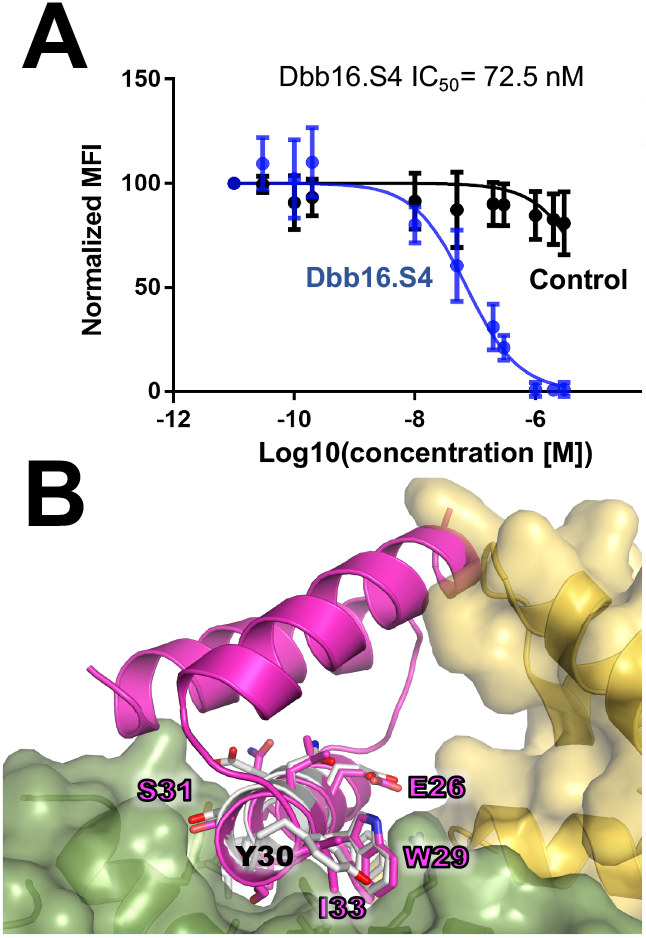
In vitro inhibition and proposed mode of action. (A) Inhibition of DBP-II binding to reticulocytes. (B) Model of the inhibitor (pink) bound to one unit of DBP-II (green) which avoids binding to the host receptor DARC (helical interface fragment in grey); the inhibitor would clash with the second DBP-II domain (yellow) and thereby would prevent their dimerization.

## Discussion

Malaria affects a third of the world’s populations and new means to treat and diagnose this disease are necessary, especially for *P. vivax*. Here, we first generated a database of highly stable three helical bundles (3H) as new protein scaffolds that can endure high concentration of protease and have high melting temperatures. These proteins can serve as a starting point for both traditional engineering projects as well as scaffolds for the computational redesign of new protein binders or inhibitors. We further were able to reveal basic parameters that determine minimum thresholds for the 3H bundles in order to be folded. Secondly, we proceeded to utilize this resource to develop a receptor mimicry to target the DBP-II protein of *P. vivax*. DBP-II is displayed during its merozoite stage and binds to ectodomain of DARC as a critical step for reticulocyte invasion of the parasite (*9, 10, 16–18*). We grafted the helical interface of DARC into several hundreds of these starting scaffolds and redesigned for optimized intra- and interface contacts with DBP-II. Designed models to test experimentally were then filtered based on identified thresholds, computed binding energy and *in silico* refolding evaluations. Six designs were monodispersed and folded proteins out of which we optimized binding affinity for the smallest protein which also introduced a clash with the backbone of the second DBP-II domain and thereby could potentially inhibit dimerization.

The lead inhibitor designed and optimized here exhibits single digit nanomolar binding affinity to DBP-II and thereby should effectively disrupt the interaction between DBP-II and DARC and hinder DBP dimerization. Based on our model of the designed protein, a dual mode of action is expected: first blocking of the DBP-II:DARC binding pocket and second preventing homo-dimerization of DBP. The designed protein inhibitor is highly resistant to protease digestions and can endure high temperatures necessary for stability in the body and for storage and distribution. Strikingly, the designed inhibitor readily refolds upon denaturation and assumes it original conformation. We thereby believe our designed and optimized inhibitor could be a valuable therapeutic candidate for treatment of the disease.

## Materials and Methods

### De novo helix design protocol

Each protein secondary structure can be described as a sequence describing the bins of the *phi-psi* angles in the Ramachandran plot; which we categorize into 5 different bins termed ABEGO(*37*). To build the tertiary structure of a protein, short structured fragments with the desired ABEGO sequence are used for its assembly. Hence, previously, the tertiary structure to be built was recorded as a single ABEGO sequence in an individual blueprint file. We generated blueprints for all possible permutation of two groups of helix ranges (group 1: 14, 15, 16 residues; and group 2: 17, 18, 19 residues) together with four types of different loop conformations (BB, GBB, GB, GABB). Blueprints were generated using a custom python script (data files). Using the same computing power (2.1 mHz Intel Xeon) and filters, we thereby allowed “scouting” runs of 432 design trajectories (one for each blueprint) producing a range of possible backbones. Since filters were applied, not all of them produced output.We then used the most “productive” blueprints (blueprints that produced more than 10 designs in 2 h) to generate more models.

The design protocol starts by building a backbone in the centroid mode (*37*) through fragment insertion with desired secondary structure conformations. As simplified scoring function we used scoring terms cenpack, hbond_sr_bb, hbond_lr_bb, atom_pair_constraint, angle_constraint, dihedral_constraint. If the backbone had the desired secondary structure conformation, and helices not bend past 15 degrees, sequence design was initiatied and attempted 10 times in order to fulfill the following criteria: average degree of at least 9 amino acids within a radius of 10 Å, less than 6 alanine residues and a packing score above 0.58. Obtained decoys were then filtered in a second script by fa_atr_per_res, local four-mer geometry, cavities, contact_per_residue and mismatch of local structure with predicted secondary structure based on Psipred (*37*); see filter_HHH.xml protocol for details. Resulting models were then evaluated through *ab initio* structure prediction to confirm that they were at their global energetic minimum using a modified protocol with reduced numbers of fragments (*38*) for the prediction process.

#### Design of DBP binding proteins

Using the most recent implementation of Epigraft^16^ in form of the MotifGraft mover, segments of the core interface helix of DARC (residue 22 – 30) of varying lengths extracted from the co-crystal structure (PDB ID 4nuv) were aligned with each scaffold using fragment superposition, not endpoint joining. If the rms distance between the joining backbone endpoints was less than 0.25 Å and diverged less than 1.0 Å overall when aligning the replaced scaffold fragment with the interface helix of DARC, grafting was attempted. All predicted interface residues of suitable scaffolds were substituted with alanine and, using backbone minimization, we attempted to close broken bonds. Then both contacts interfacing with the grafted element as well as all residues of the grafted helix except for residues 22, 23, 25, 26, 28, 29, 30 (which were declared as “hotspots” within the grafting mover) were redesigned. Both intra- and intermolecular interfaces were then extensively redesigned using 3 rounds of the most current RosettaDesign^20^ while also allowing rigid-body minimization. The best scoring design for each successful graft solution was kept if the computed binding energy below −24 Rosetta energy units (REU). Designs with a lower then −3 REU per residue score, a higher degree of connectivity of 10 were subjected to forward folding and designs that appeared at a clear global minimum were ordered for experimental evaluation.

#### Protein expression and purification of designed proteins

Amino acid sequences of designed proteins were encoded into DNA using DNAworks2.0 and “ecoli2” codons(*30*). Genes encoding the proteins were cloned into pET29b between the restriction sites *NdeI* and *XhoI* and expressed for 4 h in Terrific Broth (BD Difco) using Lemo21 (NEB) cells at 18°C using IPTG 0.5 mM. Cells were re-suspended in 35 ml phosphate buffered saline (PBS, 150 mM NaCl and 25 mM phosphate buffer at pH 7.4) and lysed using a M110P Microfluidizer or a sonicator. Insoluble cell debris was removed by centrifugation for 20 min at 40,000 x *g*. Supernatant was applied to gravity-flow columns containing 1 mL of Ni-NTA for each 500 mL of culture, washed with 50 ml PBS and 50 ml PBS containing 30 mM imidazole. Proteins were eluted with 20 ml of 250 mM imidazole in PBS., followed by Ni-NTA (QIAgen) purification. The elution fraction was subjected to further cleaning *via* SEC.

#### Library Generation and Yeast Surface Selections and Titrations

Genes were cloned into pETCON^23^ and expressed as Aga2 fusions and screened by fluorescence activated cell sorting^18^. Preparation for deep-sequencing using a MiSeq(Illumina) was performed as previously described^24^.

#### Protease digestion of scaffold library

Protease reagent Trypsin-EDTA (0.25%) solution was purchased from Life Technologies and stored at stock concentration (2.5 mg/mL) at −20°C. α-Chymotrypsin from bovine pancreas was purchased from Sigma-Aldrich as lyophilized powder and stored at 1 mg/mL in Tris buffered saline (20 mM Tris, 100 mM NaCl, pH 8.0) (TBS) with 100 mM CaCl2 at −20°C. Each reaction used a freshly thawed aliquot of protease reagent. EBY100 yeast cell cultures were induced in SGCAA for 16-18 h at 22°C for the first round and 30°C for the second round of sorting (*39*). For each protease condition, 1-3×10^8^ cells were aliquoted, pelleted, and washed in 1.00 mL of TBS. Proteolysis was initiated by re-suspending the pelleted cells in 1.00 mL of room temperature protease mix at 3 different protease conditions: 0.05 μM trypsin and 0.01 μM chymotrypsin (“Low”); 0.25 μM trypsin and 0.05 μM chymotrypsin (“Medium”); and 1.25 μM trypsin and 0.25 μM chymotrypsin (“High”). After 5 min of incubation at room temperature, cells were pelleted for 1 min and washed with 1.00 mL of chilled PBSF (20 mM NaPi, 150 mM NaCl, pH 7.4 (PBS) with 1% bovine serum albumin (BSA)); this was repeated once more to wash out the protease.

#### Library selections for binding to biotinylated DBP-II

##### Library Preparation and Next-Generation Sequencing

Plasmids were extracted from 5×10^7^ yeast cells for each selection and starting pool as previously described(*40, 41*). After *ExoI* (NEB) and Lambda exonuclease (NEB) treatment, the Illumina sequencing primers sequence, flow cell adapters and selection specific-barcodes were added *via* 2 nested PCRs using NEBnext. They additionally add 12 entirely degenerate bases at the beginning of the forward and reverse read, ensuring adequate diversity for the Illumina base-calling algorithms. A 250-PE kit was sequenced with 15% phiX at 10 pM concentration using a MiSeq (Illumina). After quality control filtering(*40*), we calculated enrichment for each sort by dividing the frequencies of selected by ones seen in the starting pool.

##### Yeast surface display (includes protease)

For library transformation, EBY100 (*39*) cells were transformed with 2 μg of the gene library together with 1 μg linearized pETCON (*SalI, XhoI, NheI* digested)(*42*) using previously reported protocol (*43*). Transformation resulted in 2-5×10^7^ cells. Yeast cells were induced for about 16-18 h at 23°C (protase sorts of *de novo* scaffolds) or 30°C, then washed once with PBSF. Protease-digested cells were labeled with 10 ng/ml anti-c-Myc antibody conjugated to FITC for ≥15 min on a rotator at room temperature. For the first round of selections of the SSM library, cells were incubated for 2 h at 23°C with 1 μM bDBP-II, before adding a fourth of the used concentration of streptavidin conjugated to phycoerythrin (SAPE, Invitrogen) and 2 ng/ml anti-c-Myc antibody conjugated to FITC and incubating for another hour. For non-avid conditions in later sorts, cells were incubated for 1 h at 23°C while rotating, unless noted otherwise, and then washed once with ice-cold PBSF before re-suspending in 100 μL with 72 nM SAPE and 2 ng/ml anti-c-Myc antibody conjugated to FITC for 15-20 min on ice. Cells were washed once with 1 mL ice-cold PBSF and stored as pellets on ice before their final re-suspension in 1 mL ice-cold PBSF followed by fluorescence-activated cell sorting (FACS).

##### Library Preparation and Next-Generation Sequencing

Plasmids were extracted from 5×10^7^ yeast cells for each selection and starting pool, and 6.7×10^7^-1.3×10^8^ yeast cells from the protease-digestion sorts, as previously described(*40, 41*). After *ExoI* (NEB) and Lambda exonuclease (NEB) treatment, the Illumina sequencing primers sequence, flow cell adapters and selection specific-barcodes were added *via* 2 nested PCRs using NEBnext. They additionally add 12 entirely degenerate bases at the beginning of the forward and reverse read, ensuring adequate diversity for the Illumina base-calling algorithms. A 250-PE kit was sequenced with 15% phiX at 10 pM concentration using a NextSeq (Illumina). After quality control filtering(*40*), we calculated enrichment for each sort by dividing the frequencies of selected by ones seen in the starting pool.

##### Binding analysis

Titrations were performed at 30° C while rotating at 1000 rpm on an OctetRED96 BLI system (ForteBio, Menlo Park, CA) using streptavidin-coated biosensors. Sensors were equilibrated for 20 min in PBSTB buffer (PBS, 0.002% Tween 20, 0.01% BSA). Each sensor was loaded with 20 nM biotinylated DBP-II for 100 sec. For baseline, association and dissociation time intervals of 60 sec, 240 and 300 sec were applied respectively.

### Protein production and biotinylation of avi-tagged DBP-II

BirA-tagged Sal-1 DBP-II was prepared as described previously (*9–11, 23*). Briefly, inclusion bodies were solubilized in 6 M guanidinium hydrochloride and refolded via rapid dilution into 400 mM L-arginine, 50 mM Tris (pH 8.0), 10 mM EDTA, 0.1 mM PMSF, 2 mM reduced glutathione, and 0.2 mM oxidized glutathione. Refolded protein was captured on SP Sepharose Fast Flow resin (GE Healthcare) and further purified by size-exclusion chromatography (GF200; GE Healthcare) into 10 mM Hepes (pH 7.4) and 100 mM NaCl.

BirA-tagged Sal-1 DBP-II was buffer exchanged into PBS. Then 50 μL of BiomixA (Avidity), 50 μL of BiomixB (Avidity), and 10 μL of 5 mM *D*-biotin (Avidity) were added to the protein along with BirA ligase, followed by overnight incubation at 4 °C. The biotinylation was confirmed by Western Blot using Streptavidin HRP-conjugate (Thermo Scientific). Before use, the reaction mix was buffer-exchanged into PBS.

### Fluorescence Activated Cell Sorting Assay

Biotinylated purified DBP-II protein in concentration of 10 nM was incubated with eleven different concentrations of Dbb 16T (from 0.03 nM to 1 μM interval of concentrations) for 1 h at RT. Red blood cells were added and incubated for an additional 1 h at RT. To detect bound DBP Alexa Fluor 488 Streptavidin conjugate (Fisher) was added to the mix and incubated at RT for 1 h followed by washing twice with PBS. FITC labeling was measured by flow cytometry. An IC50 value for Dbb 16T was calculated in GraphPad Prism from five independent biological replicates. IC50 curve for the coil peptide of similar size was plotted as a negative control.

## Author contributions

EMS and NHT conceived the study. ART, RC, DU, JCK and EMS performed experiments. EMS designed all proteins. ART, DU, NHT and EMS analyzed the data.

EMS and NHT wrote the manuscript. All authors read and commented on the manuscript.

## Data and materials availability

detailed protocols are either attached as XML files or described under the supplementary information section.

